# Adaptive adjustment of significance thresholds produces large gains in microbial gene annotations and metabolic insights

**DOI:** 10.1101/2024.07.03.601779

**Authors:** Kathryn Kananen, Iva Veseli, Christian J. Quiles Pérez, Samuel Miller, A. Murat Eren, Patrick H. Bradley

## Abstract

Gene function annotations enable microbial ecologists to make inferences about metabolic potential from genomes and metagenomes. However, even tools that use the same database and general approach can differ markedly in the annotations they recover. We compare three popular methods for identifying KEGG Orthologs, applying them to genomes drawn from a range of bacterial families that occupy different host-associated and free-living biomes. Our results show that by adaptively tuning sequence similarity thresholds, sensitivity can be substantially improved while maintaining accuracy. We observe the largest improvements when few reference sequences exist for a given protein family, and when annotating genomes from non-model organisms (such as gut-dwelling Lachnospiraceae). Our results suggest that straightforward heuristic adjustments can broadly improve microbial metabolic predictions.

## Introduction

Metabolism, which links biochemical transformations to the survival and proliferation of organisms, is key to understanding microbial ecology and evolution. One important way to derive insights about microbial metabolism at scale is through (meta)genome annotation. Modern annotation tools can leverage decades of labor-intensive, mechanistic work on gene function to make high-throughput *de novo* inferences about metabolic potential. However, some of the most accurate methods, which use phylogenetic information and/or predictions of protein structure to find orthologs of genes of known function, are computationally demanding and scale poorly to increasingly large ‘omics surveys. Sequence homology searches offer a more scalable and streamlined approach to annotation, but even when the same reference databases and alignment methods are used, implementation details can affect the rate of gene function recovery.

To systematically assess how annotation strategies affect annotation quantity and quality, we annotated KEGG Orthologs (KOs) [1] in 396 genomes from 11 bacterial families, representing five different biomes and ranging from clades that included common model organisms and pathogens (*Enterobacteriaceae*) to less-well-studied families (*Lachnospiraceae, Nanosynbacteriaceae*). We used three methods that all rely on the KOfam database of profile hidden Markov models (pHMMs) but process alignment scores differently to produce annotations: Kofamscan [2], MicrobeAnnotator [3], and anvi’o [4]. We then used these annotations to estimate metabolic pathway completeness.

## Results

All three tools begin by aligning ORFs to KOfam pHMMs and then applying a bit-score threshold. For all pHMMs based on at least three sequences, KOfam provides such a threshold derived from cross-validation. The three methods then diverge in two main ways. First, some ORFs have no matches above the cutoff: Kofamscan by default discards these. In contrast, to catch what we call “weak homologs,” anvi’o applies a second filter where the threshold is lowered by 50% (by default), then tests whether all matches passing both this filter and global e-value threshold are from the same KOfam (Supplementary Figure 3). MicrobeAnnotator can annotate weak homologs by BLASTing ORFs against a second database (SwissProt), transferring top KO annotations above a global threshold. Second, some KOfams lack a cross-validation threshold (we call these “nt-KOs”, short for “no threshold”). By default, neither Kofamscan nor anvi’o attempts to annotate nt-KOs. However, anvi’o can optionally apply a conservative adaptive threshold, based on the lowest bit-score from sequences used to build the pHMM for each nt-KO. MicrobeAnnotator takes a more permissive approach, treating any match to nt-KOs as legitimate, regardless of bit-score or e-value. A full explanation can be found in Supplementary Methods.

We observed large differences in annotation rate between methods. MicrobeAnnotator and anvi’o, which employ additional heuristics for recovering homologs, consistently annotated more ORFs per genome compared to Kofamscan (Figure 1a). Using the bit-score heuristic in anvi’o enabled recovery of ∼13% more annotated genes per genome (range: 3.1% to 21%, paired Wilcox test p<2.2×10^−16^). Disabling this heuristic made the results identical to Kofamscan’s. To assess annotation quality, we compared to an orthology-based method, eggNOG-mapper, which transfers annotations via similarity to a fine-grained pre-computed orthologs database (eggNOG v5) [5] using DIAMOND [6]. Because it relies on a curated database, we expected highly accurate but possibly less complete results. Anvi’o returned the largest percentage of KO annotations across all genomes confirmed by eggNOG-mapper, with 42% (608,311/1,436,238 KOs), followed by MicrobeAnnotator at 40% (577,041/1,436,238 KOs). Kofamscan recovered the smallest number (576,853/1,436,238 KOs, 40%). Comparing total annotations recovered across all genomes, MicrobeAnnotator and anvi’o had nearly the same percent increase of 13% more annotations recovered than Kofamscan. However, when only confirmed annotations are considered, 4.4% more genes were recovered per genome on average using anvi’o (range: 1.1% to 15%, p<2.2×10^−16^). We also ran MicrobeAnnotator in “refine” mode, which additionally maps genes against SwissProt and converts Enzyme Commission (EC) numbers and InterPro accessions to KO accessions; however, additional KOs recovered were not unique and ultimately resulted in approximately 0% more annotations per genome.

**Figure 1.**
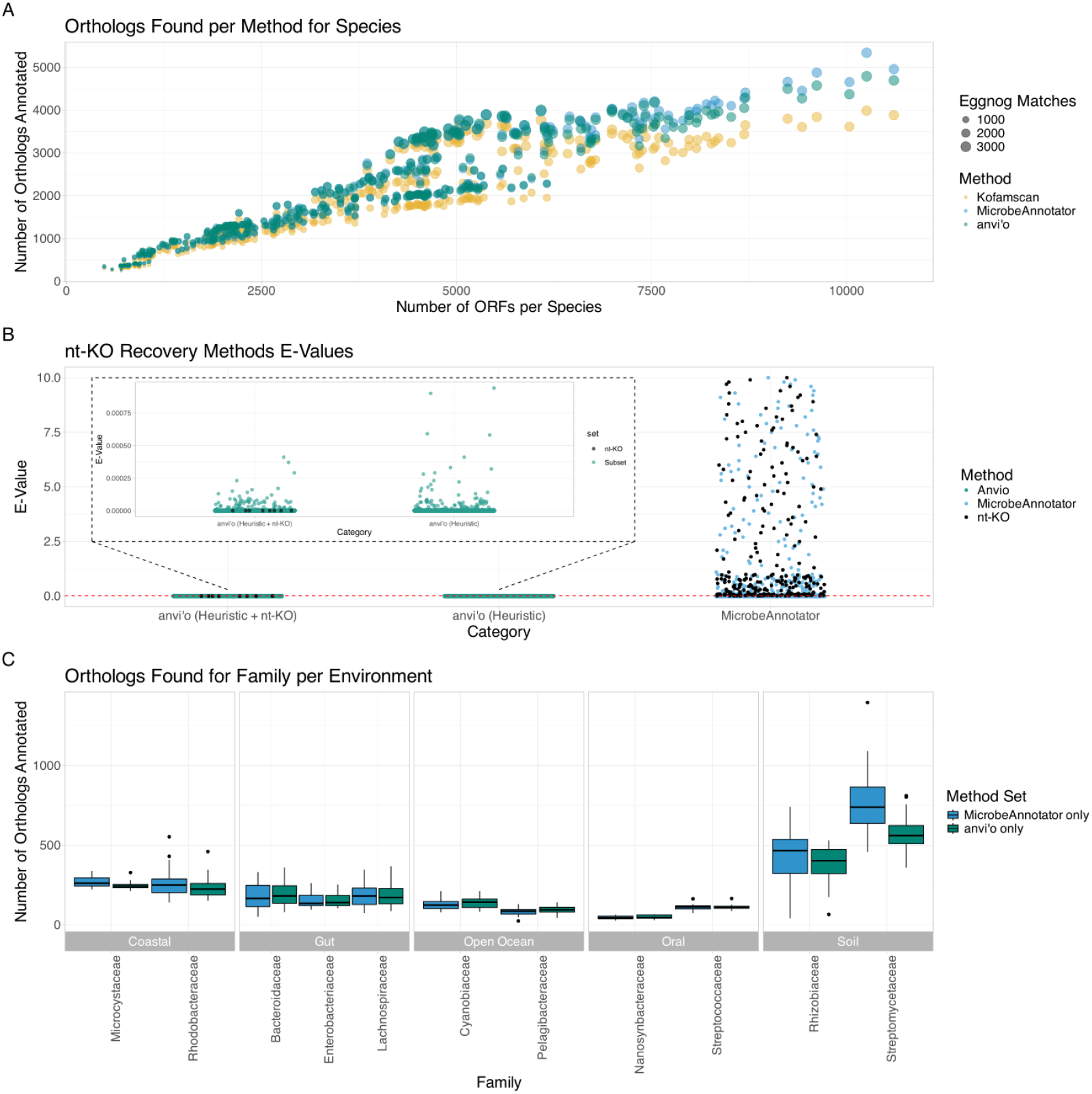
Heuristics consistently improve annotation rate. **A**. Comparison of the total KOs identified by each tool with default parameters. The x-axis displays the number of open-reading frames (ORFs) in the genome. The size of each point corresponds to the number of orthologs that were also found by EggNOG-mapper 5.0 using the EggNOG database as further evidence of correct KO annotation. **B**. Distribution of e-values for KOs and nt-KOs as annotated by anvi’o or MicrobeAnnotator. The plot includes all unique e-value scores from each respective set of KO annotations from anvi’o run with default parameters, anvi’o run with the ‘--include-stray-KOs’ flag, and MicrobeAnnotator run with default parameters. The black points identify nt-KOs. The standard e-value threshold of 0.01 is indicated with the dashed red line. **C**. KOs that are uniquely identified by only one annotation method in each microbial family. Very few orthologs were identified by both anvi’o and MicrobeAnnotator but not Kofamscan (median was 1 per family).

Because MicrobeAnnotator always annotates nt-KOs, we ran an additional comparison enabling nt-KO annotations in anvi’o. This increased its annotation rate on average by 0.0705% per genome (p<2.2×10^−16^), yielding approximately 1,536 ORFs with confirmed annotations, compared to 1,457 from MicrobeAnnotator. Notably, for MicrobeAnnotator, annotation e-values repeatedly fell outside the standard cutoff of e<0.01, with many surpassing 1 (Figure 1b). Thus, nt-KOs annotated using anvi’o are more likely to be reliable.

To determine how annotation rates varied across bacterial taxa, we counted KOs annotated uniquely by MicrobeAnnotator or anvi’o (none were uniquely annotated by Kofamscan). Between 24 and 1,396 orthologs were uniquely annotated by MicrobeAnnotator, and between 28 and 811 with anvi’o (Figure 1c). *Enterobacteriaceae* genomes had a lower number of unique KOs, likely because this family contains *E. coli* along with several common pathogens, and is expected to be well-represented in KEGG. In contrast, many unique annotations were recovered for *Rhizobiaceae*, a family containing numerous plant-associated microbes with only 7K genomes in GTDB, compared to 97K for *Enterobacteriaceae*. All annotations unique to MicrobeAnnotator were nt-KOs.

To assess downstream impacts of variable annotation recovery, we examined metabolic module completeness scores for all 11 families. Using anvi’o, we calculated KEGG module completeness [7] from each tool’s default annotations. For each family, we also determined the 10 modules whose completeness score differed most across tools (Supplementary Table 3). Overall, anvi’o resulted in 11% more modules per genome with ≥80% completeness compared to MicrobeAnnotator, and 12% more compared to Kofamscan. Averaging across all genomes in a family, median completeness scores increased in 15% of modules with anvi’o and in 13% with MicrobeAnnotator. For anvi’o, median completeness scores improved most often for less-well-studied families, such as *Nanosynbacteraceae* (higher than Kofamscan in 38% of cases), and least often for the well-studied *Enterobacteriaceae* (higher in 4.9% of cases). MicrobeAnnotator did not display this trend (Supplementary Table 3, Supplementary Figure 1). Furthermore, at ≥80% completeness, anvi’o uniquely identified 142 modules at the species level (compared to 1,544 modules identified by all tools). We did identify modules unique to MicrobeAnnotator, especially drug resistance pathways (Figure 2b, Supplementary Figure 2). However, these appeared unlikely given biological context: annotations from MicrobeAnnotator suggest a complete carbapenem resistance pathway (M00851) in all families except *Synechococcaceae* and *Nanosynbacteraceae*, whereas anvi’o and Kofamscan only detect this in *Enterobacteriaceae*. While carbapenem resistance was previously reported in some marine microbiota [8], to be found fully complete among all species in nearly all families is unlikely. Instead, these annotations were likely driven by MicrobeAnnotator’s lenient approach to identifying nt-KOs.

**Figure 2.**
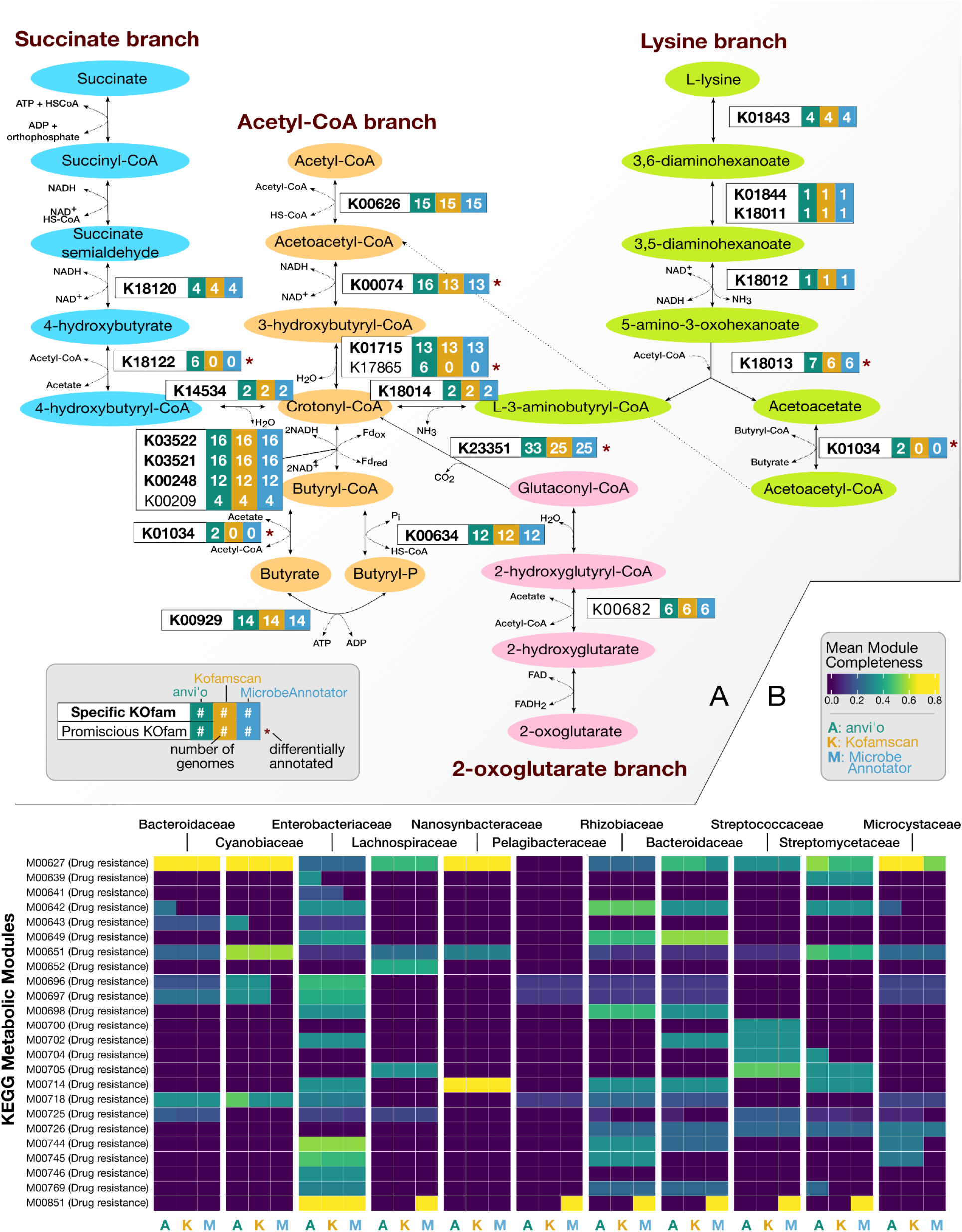
**A**. Annotated butyrate production pathway, demonstrating the number of genomes with annotation(s) for each enzyme from each of the three software tools (with default parameters). Pathway diagram is based on [24]. Substrate-specific enzymes have bold labels, while those that are promiscuous are labeled in regular text. Asterisks indicate enzymes for which at least one of the three tools yielded different results. **B**. Per-family mean completeness scores for all drug resistance modules in KEGG recovered by each method.

As a specific test case, we next compared each tool’s ability to annotate enzymes involved in biosynthesis of butyrate, a key metabolic product of gut microbes. This pathway, previously found to be complete in about 41% of *Lachnospiraceae* genomes, can terminate in two branches: butyryl-CoA transferase (BCoAT) or butyryl kinase (buk) [9]. We curated a gene set for clostridial butyrate production, then computed completeness in 36 *Lachnospiraceae* genomes. While all three tools comparably annotated the trunk of the pathway from acetyl-CoA (Supplementary Figure 4, Supplementary Table 2), anvi’o detected the KOfam for the second step (3-hydroxybutyryl-CoA dehydrogenase) in three additional genomes. Anvi’o also detected the BCoAT branch in two genomes, while the other tools found it in none (Figure 2a). Furthermore, anvi’o exclusively detected two additional KOfams (K17865, K18122) in six genomes each, and identified a glutaconyl-CoA decarboxylase with predicted involvement in butyrate production from amino acids and/or α-ketoglutarate in almost all *Lachnospiraceae* (33/36, vs. 25/36 for Kofamscan and MicrobeAnnotator). With anvi’o, this pathway was >80% complete with at least 5/6 of core operon genes detectable in 33.3% of these genomes, consistent with previous estimates [9].

While many microbial genes will remain “true unknowns”, requiring sophisticated computational and experimental efforts to determine their function, our results suggest hundreds of genes in bacterial genomes may be overlooked by standard functional annotation approaches, with small tweaks leading to substantially higher recall. In particular, moderate and adaptive relaxation of bit-score thresholds defined for KOfam models implemented in anvi’o improves annotation rates without introducing many false positives for weaker homologs. Paired with better recovery of gene families with few existing representative sequences (here, nt-KOs), we can identify more metabolic pathways with higher completeness. Reevaluating existing annotations using this heuristic may be beneficial, especially for organisms not well represented in public databases.

## Code Availability

Scripts and workflow are available at https://github.com/pbradleylab/2023-anvio-comparison.

## Acknowledgments

This work was supported by NSF grant OCE-2019589 to the Center for Chemical Currencies of a Microbial Planet and NIH grant R35GM151155 to PHB. This is C-CoMP publication #042.

## Notes

https://github.com/pbradleylab/2023-anvio-comparison

### Competing Interest Statement

The authors have declared no competing interest.

