## Supplementary Information and Supplementary Figures 1-5 for "Adaptive adjustment of significance thresholds produces large gains in microbial gene annotations and metabolic insights"

### Supplementary Info

#### Detailed methodology of benchmarked software

MicrobeAnnotator [3], Kofamscan [2], and the `anvi-run-kegg-kofams` command in anvi'o [4] are programs that annotate KOs by querying the KOfam database via the HMMER software [10]. Most pHMMs in the KOfam database are associated with a predefined bit-score threshold that these tools use to differentiate between strong and weak matches to the model. That is, when a query gene sequence has some homology to a KOfam model, the sequence is considered homologous enough for annotation when the match's bit-score is at or above the model's bit-score predefined threshold.

The primary difference between these tools' annotation methods lies in their strategy for enabling the annotation of weak homologs to KEGG Orthologs. MicrobeAnnotator addresses this problem by the use of multiple reference databases beyond KEGG and enabling the conversion of annotations from these databases into KOs (via the `--refine` parameter). Kofamscan, on the other hand, implements an optional parameter for the relaxation of bit-score thresholds (`--threshold-scale`) that applies globally (i.e., across all KOfam models), and additionally enables expectation value thresholding via the `--e-value` parameter. The anvi'o program `anvi-run-kegg-kofams` takes a more nuanced approach to relaxing bit-score thresholds: it re-visits genes that are not annotated using the predefined bit-score thresholds and assesses the set of HMM hits to these genes with bit-scores above a given fraction of the KOfam model's original bit-score threshold (default fraction is 0.5) and e-values below a given e-value threshold (by default, 1e-05). If all of these slightly-less homologous hits are from a unique KO model, then the gene is annotated with that KO (Supplementary Figure 3).

Some KOfam models (henceforth referred to as 'nt-KOs' to stand for 'no-threshold KOs') lack a bit-score threshold for differentiating between strong and weak hits because they are built from too few sequences to use the threshold estimation workflow described in Aramaki et al. [2], and the three tools also differ in how they handle annotations to these gene families. MicrobeAnnotator annotates any gene with a match to one of these nt-KOs regardless of the match's similarity score. Depending on the requested output format, Kofamscan either does not annotate these nt-KOs at all or adds an asterisk beside the annotation in the output file to indicate its potentially dubious nature. Anvi'o by default does not use these KOfam models but allows users to include them in its annotation workflow via the `--include-stray-KOs` parameter, which relies upon a conservative bit-score threshold computed within anvi'o for each of these models by taking the minimum bit-score of matches between the model and the gene sequences used to create it. These approaches differ in their potential to introduce false positive annotations via the inclusion of poor matches to this subset of KOfam models without predefined bit-score thresholds.

#### Additional comparison

When we set Kofamscan's global parameters `--threshold-scale` and `--e-value` to 0.5 and 1e-05, respectively, to match the settings used by anvi'o's bit-score heuristic, this 'refined' mode recovered an additional 35.43% more annotated genes per genome, on average. However, this increase drops to 9.99% when only annotations confirmed by EggNOG-mapper are considered.

#### Methods

##### 0. Databases

GTDB v214 [11] was used for genome data and taxonomic classification. Databases used by methods include KOfam (downloaded on 2024-03-26) [2], UniProt Swissprot (2024-03-26) [12], RefSeq (2024-03-26) [13], and Trembl (2024-03-26) [14]. EggNOG-mapper v5.0 [5, 15] and its accompanying database were used for the independent validation of annotations.

##### 1. Resources and Environment

Mambaforge v1.5.8 was used for package management. Snakemake v7.30.1 [16] was used for workflow management. Eido v0.2.2 and Peppy v0.35.7 [17] were used to generate sample input tables and run PEP. High-performance computing resources were supplied by the Arts and Science College, The Ohio State University.

##### 2. Sample Selection

Potential genomes were randomly subsampled without replacement (n=36) per selected family from GTDB for 11 families from different biomes [11, 18–20]. Only genomes with publicly available annotations were retained. Files were retrieved with ncbi-genome-download v0.3.3 [21].

##### 3. Gene Calling

Prodigal v2.6.3 [22] was used for gene calling for all methods.

##### 4. Protein Translation

Anvi'o v8-dev (commit 545ba63) [4] was used to generate protein fasta per genome.

##### 5. Kofamscan

Kofamscan v1.3.0 [2] was run twice, once using default settings and once in 'refined' mode with a threshold scale of 0.5 and an e-value threshold of $1x10^{-5}$.

##### 6. MicrobeAnnotator

Internal databases were retrieved and indexed using MicrobeAnnotator v2.05 [3] (commit 9bbc5f6). Then they were downloaded and indexed using built-in scripts. Finally, KO prediction was performed twice. Once using default settings, and then run with the `--refine` flag.

#### 7. Anvi'o

Anvi'o v8-dev (commit 0e01dd7) [4] was initially run using default settings, which include the use of the adaptive bit-score heuristic. Then it was run with the heuristic turned off, using the `--skip-bitscore-heuristic` flag. Finally, anvi'o was run with the `--include-stray-KOs` flag to incorporate annotations for nt-KOs.

#### 8. nt-KO Evaluation

Any KOs reported without an associated bit-score threshold in the KOfam database were considered nt-KOs.

#### 9. Validation of annotations

All genomes were run through EggNOG-mapper 5.0 [5, 15] and the resulting KO predictions were compared to KOs recovered from each method. Annotations that matched those found with EggNOG-mapper were considered to be validated. Unvalidated annotations include potential false positives as well as KOs that were not found in the EggNOG database.

#### 10. Pathway Analysis

In-house scripts were used to generate enzyme-txt files (https://anvio.org/help/8/artifacts/enzymes-txt/) to use as input for the anvi-estimate-metabolism program in anvi'o (https://anvio.org/help/8/programs/anvi-estimate-metabolism/) for each tool/parameter set combination. For anvi'o and Kofamscan annotations, we included in the enzyme-txt files only the hit with the lowest e-value for each gene to match the behavior of MicrobeAnnotator. Pathwise completeness scores for each metabolic pathway in the KEGG MODULE database were generated with the anvi-estimate-metabolism program. Individual pathway completeness scores were averaged across all genomes in each family (see Figure 2b, Supplementary Figures 3 and Supplementary Table 3) per method with highest number of recovered KOs (See Figure 1a). We created custom metabolic modules for butyrate biosynthesis by identifying the KOs associated with this pathway as described using Enzyme Commission (EC) numbers in [9, 23, 24], with the addition of the crotonase enzyme K17865 (EC: 4.2.1.55) for the dehydration reaction from 3-hydroxybutyryl-CoA to crotonyl-CoA. One module ('BUTANOATE') describes the full set of enzymes with all potential alternative KOs, another module ('SUBSPEC') includes only substrate-specific enzymes whenever possible, and the last module ('BUCASYNOP') includes only the 6-gene core operon for butyryl-CoA synthesis. After generating a user-defined modules database with `anvi-setup-user-modules`, we estimated pathwise completeness for these pathways for the default annotation modes of each tool within the set of 36 Lachnospiraceae genomes using `anvi-estimate-metabolism`.

#### 11. Visualization

R version v4.3.2 [25] was used to generate all graphics with the exception of Supplementary Figures 4 and 5. Barplots were generated using tidyverse v2.0.0 [26], plyr v1.8.9 [27], and ggplot2 v3.5.0 [28]. Scatterplots were generated with ggplot2. Heatmaps were generated using viridis v0.6.5 [29], data.table v1.15.4 [30], devtools v2.4.5 [31], and the ComplexHeatmap v2.15.4 package [32, 33]. In Figure 2b, drug resistance modules were plotted for the three methods by subsetting any module category that contained the string 'Drug resistance'. In Supplementary Figure 1a, pathwise-completeness anvi'o and MicrobeAnnotator over Kofamscan for each module per genome was calculated and graphed. Any modules where no change for either tool is detected were graphed (Supplementary Figure 1). For Supplementary Figure 4a, the heatmap of completeness scores was generated with the anvi'o program `anvi-interactive`. For Supplementary Figure 4b, we colored the corresponding KEGG pathway map for butanoate metabolism (accession ko00650) according to which tool(s) were able to annotate each enzyme in the map in at least one of the *Lachnospiraceae* genomes. To create the map figure, custom scripts were used to parse the annotation files, identify the set of tools annotating each KO, and use the resulting table to color the KEGG pathway map.

### Supplementary Figures

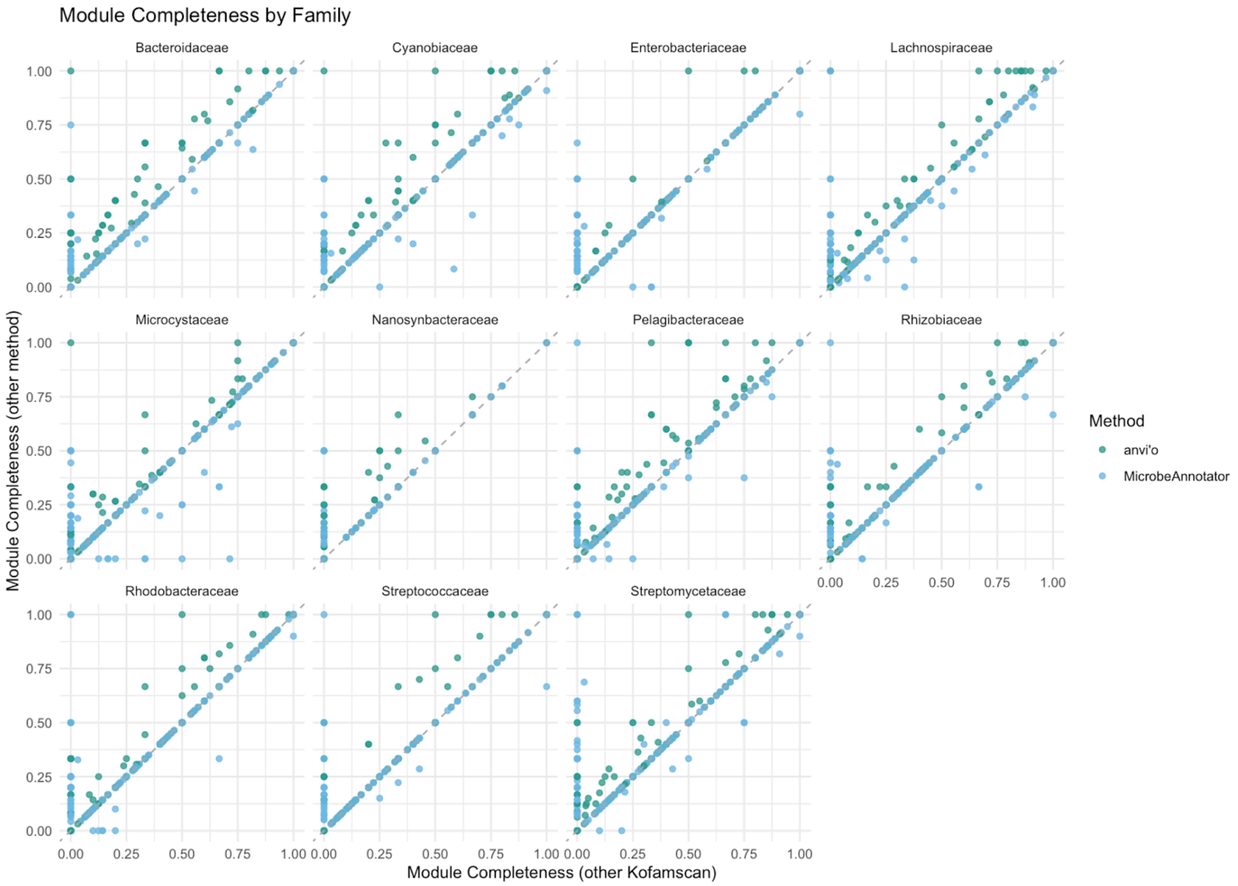


**Supplementary Figure 1.** Median module completeness of Kofamscan (x-axis) compared to anvi'o (y-axis, green) and MicrobeAnnotator (y-axis, blue). Each point is the median completeness score of one module over all genomes in a given bacterial family (facets).

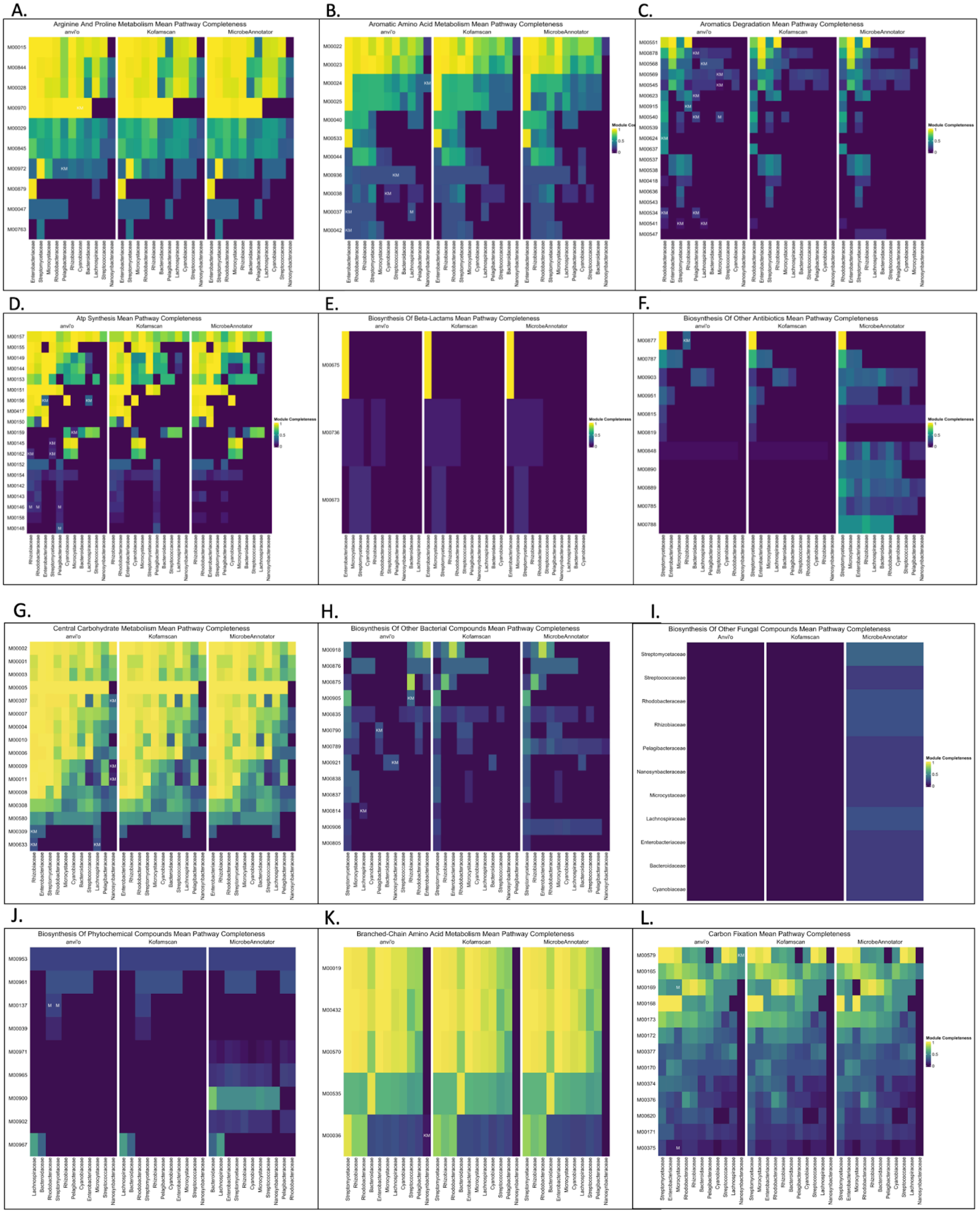
A.
Arginine And Proline Metabolism Mean Pathway Completeness
B.
Aromatic Amino Acid Metabolism Mean Pathway Completeness
C.
Aromatics Degradation Mean Pathway Completeness
D.
Atp Synthesis Mean Pathway Completeness
E.
Biosynthesis Of Beta-Lactams Mean Pathway Completeness
F.
Biosynthesis Of Other Antibiotics Mean Pathway Completeness
G.
Central Carbohydrate Metabolism Mean Pathway Completeness
H.
Biosynthesis Of Other Bacterial Compounds Mean Pathway Completeness
I.
Biosynthesis Of Other Fungal Compounds Mean Pathway Completeness
J.
Biosynthesis Of Phytochemical Compounds Mean Pathway Completeness
K.
Branched-Chain Amino Acid Metabolism Mean Pathway Completeness
L.
Carbon Fixation Mean Pathway Completeness
anvi'o
Kofamscan
MicrobeAnnotator
Module Completeness

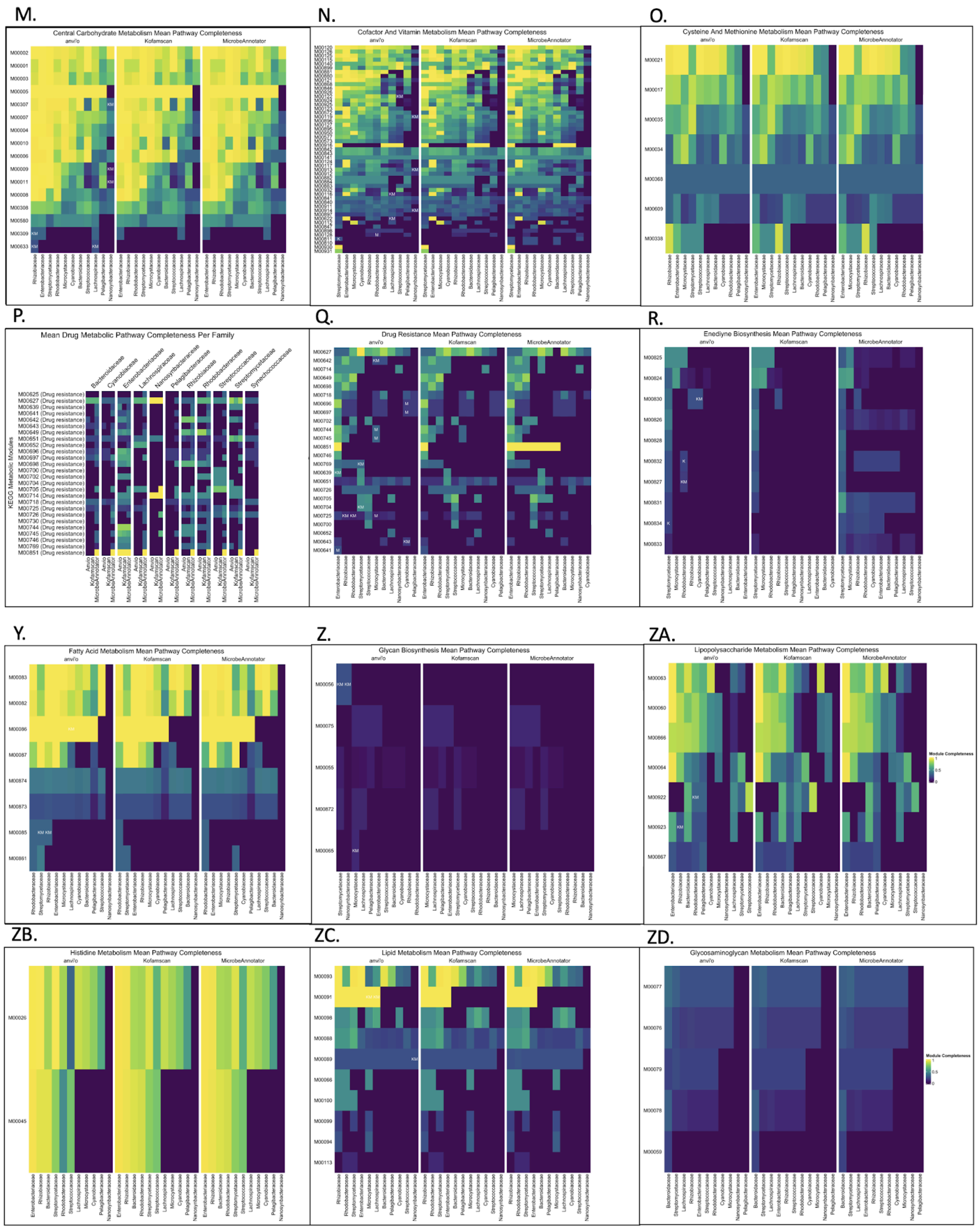
M.
Central Carbohydrate Metabolism Mean Pathway Completeness
N.
Cofactor And Vitamin Metabolism Mean Pathway Completeness
O.
Cysteine And Methionine Metabolism Mean Pathway Completeness
P.
Mean Drug Metabolic Pathway Completeness Per Family
KEGG Metabolic Modules
Q.
Drug Resistance Mean Pathway Completeness
R.
Enediyne Biosynthesis Mean Pathway Completeness
Y.
Fatty Acid Metabolism Mean Pathway Completeness
Z.
Glycan Biosynthesis Mean Pathway Completeness
ZA.
Lipopolysaccharide Metabolism Mean Pathway Completeness
Module Completeness
ZB.
Histidine Metabolism Mean Pathway Completeness
ZC.
Lipid Metabolism Mean Pathway Completeness
ZD.
Glycosaminoglycan Metabolism Mean Pathway Completeness
Module Completeness
anvi'o
Kofamscan
MicrobeAnnotator

ZE.

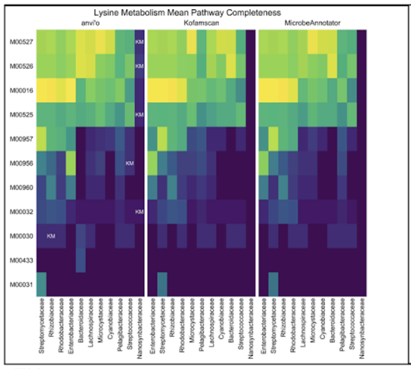


ZF.

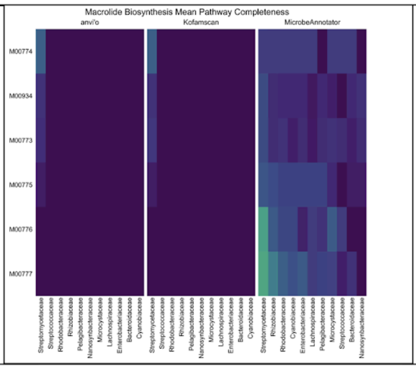


ZG.

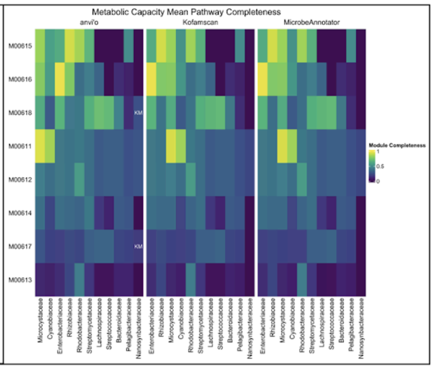


ZH.

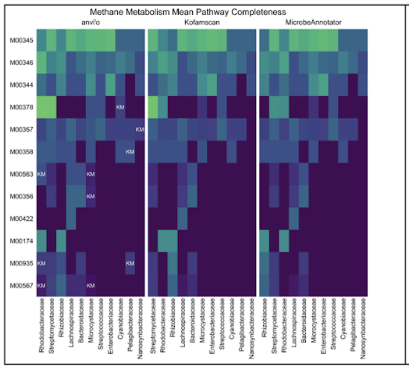


ZI.

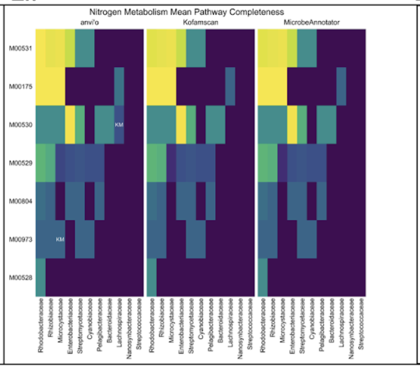


ZJ.

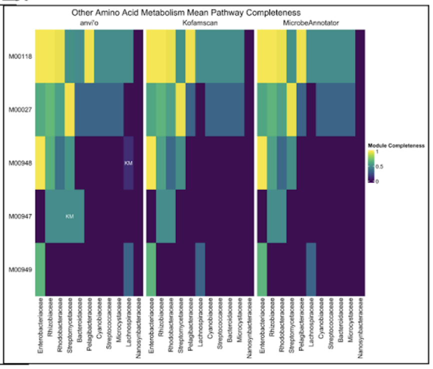


ZK.

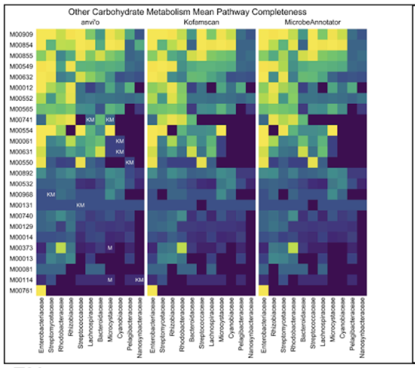


ZL.

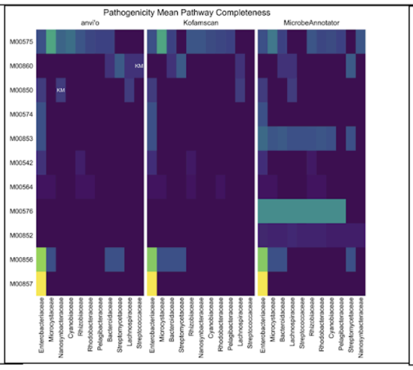


ZM.

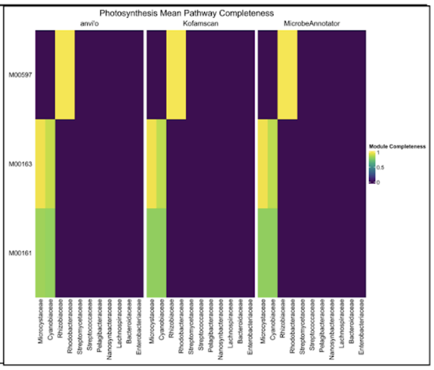


ZN.

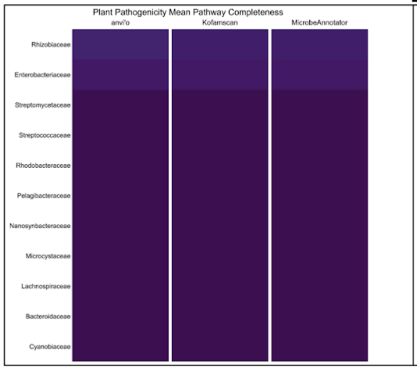


ZO.

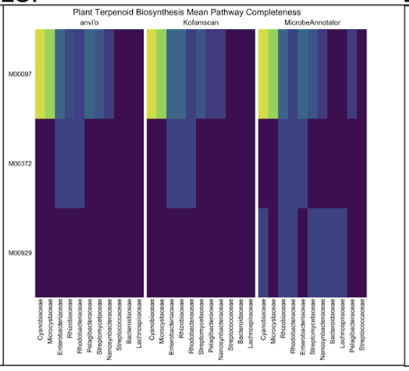


ZP.

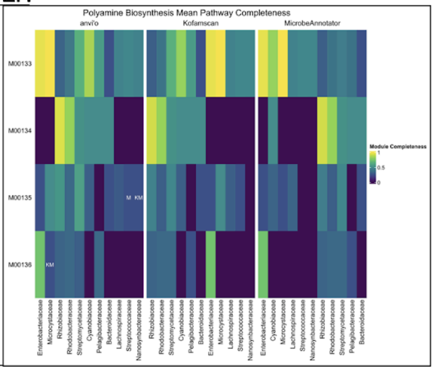

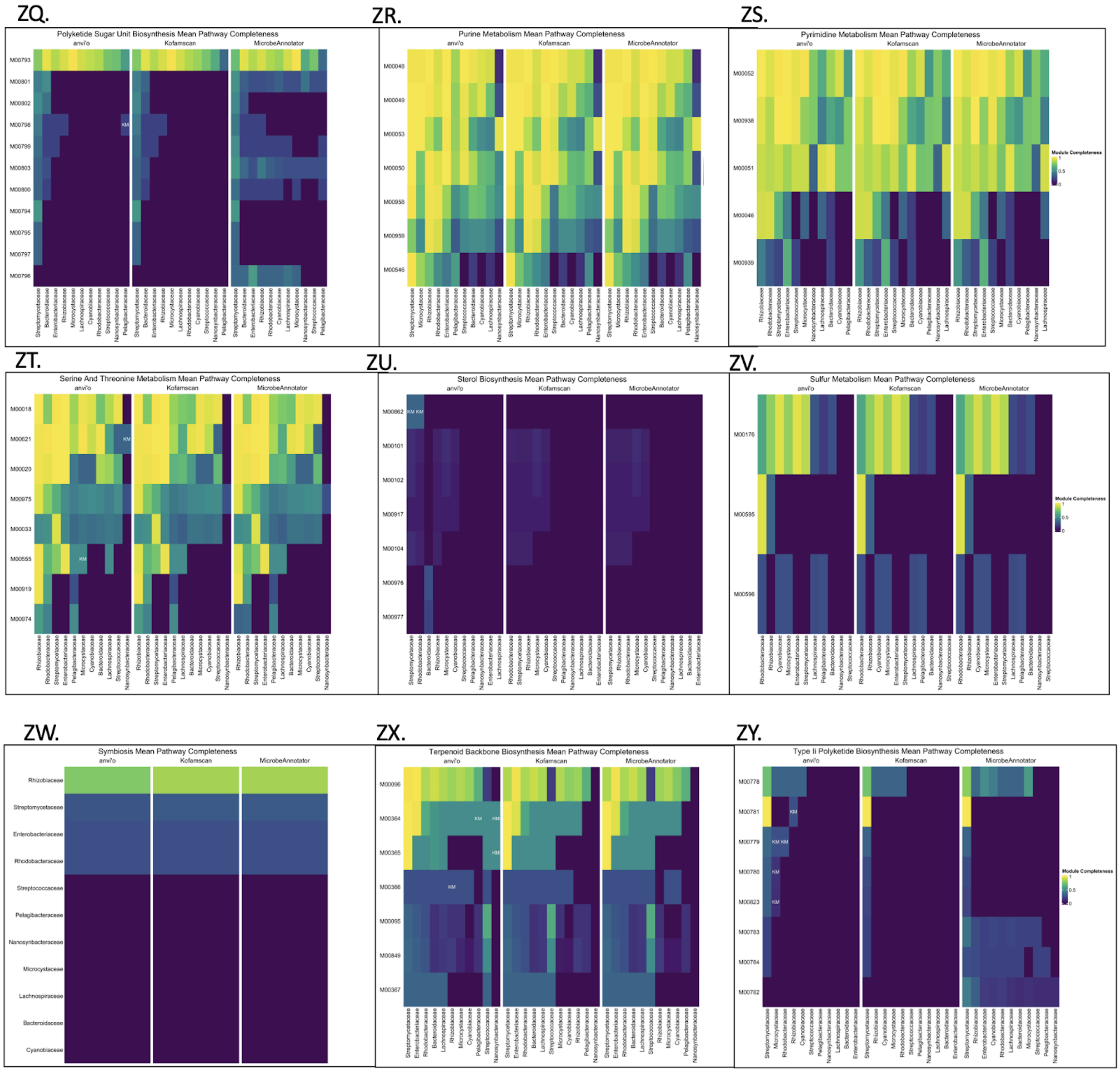


**Supplementary Figure 2.** The mean module completeness score in each microbial family for all KEGG modules found across the three methods is shown. Each module is grouped into one of 45 categories and the MicrobeAnnotator, Kofamscan, and anvi'o default results are compared. Modules predicted with >0% completeness by anvi'o are marked with an 'M' if they are not also found with MicrobeAnnotator, a 'K' if they are not found with Kofamscan, and 'KM' if they are absent (0% complete) in both of the other methods.

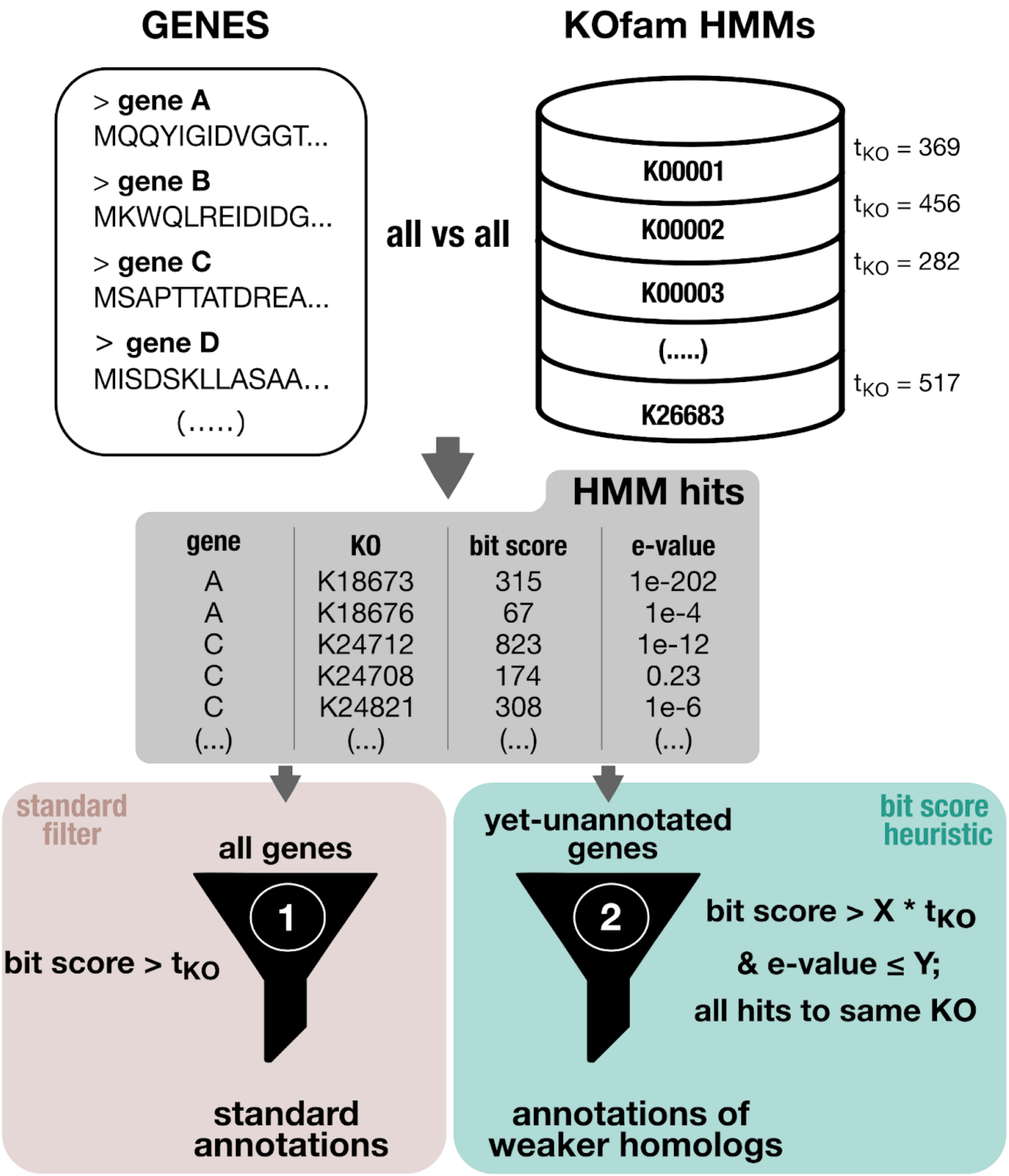


**Supplementary Figure 3. The annotation workflow and bit-score heuristic in anvi'o.** In \`anvi-run-kegg-kofams\`, HMM hits from an all vs. all query of predicted gene sequences against the KOfam database are initially filtered using the KEGG-defined bit-score threshold for each KO model, $t_{KO}$. If the bit-score heuristic is turned on (the default behavior, in green), the program does a second pass through the hits to genes that have not yet been annotated, filtering them using a percentage (X) of the original bit-score threshold and a maximum e-value cut-off (Y). If all of those filtered hits are to the same KO model for a given gene, the gene is annotated with the KO despite its slightly weaker homology to this family.

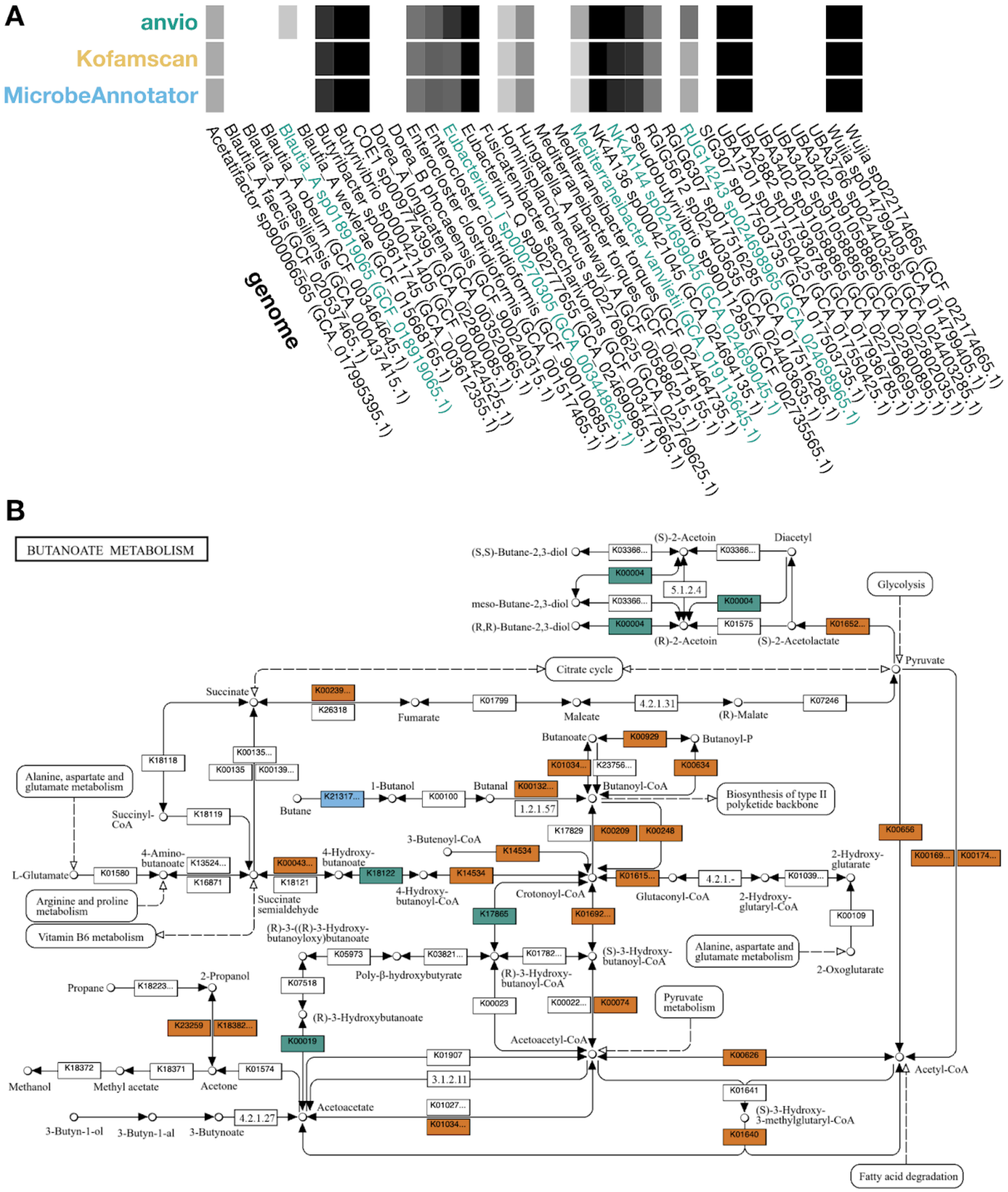


**Supplementary Figure 4.** The butyrate biosynthesis pathway as annotated in the 36 *Lachnospiraceae* genomes in our test dataset. Each tool was run with default parameters. **A.** Heatmap of pathwise completeness scores for the custom butyrate biosynthesis pathway module defined with substrate-specific enzymes whenever possible (the "SUBSPEC" module).

Genome labels are colored green when anvi'o estimates higher completeness for the pathway than the other tools. **B.** The combined annotations from all 36 genomes mapped onto the KEGG Pathway Map for Butanoate Metabolism and colored according to which tool(s) could annotate them. Enzymatic reactions are colored orange if anvi'o, MicrobeAnnotator, and KOfamscan all annotated the corresponding orthologs, green if anvi'o alone annotated orthologs, and blue if MicrobeAnnotator alone annotated orthologs. No other combinations of tools were observed in the ortholog annotations for this pathway map.

### Supplementary Tables

**Supplementary Table 1.** Describes the genomes used in our analysis and the following information for each genome: assigned taxonomic family and species, number of gene calls, number of genes annotated per method, and mean module completeness per method. The method includes both which tool was run ('kofamscan'; 'microbeannotator'; 'anvio') and with which parameters ('default': default parameters for all three tools; 'refined': non-default parameters for both Kofamscan and MicrobeAnnotator; 'noheuristic': with the `--skip-bitscore-heuristic` flag for anvi'o; 'stray': with the `--include-stray-KOs` flag for anvi'o).

**Supplementary Table 2.** Pathwise completeness scores in the 36 *Lachnospiraceae* genomes for the three custom butyrate biosynthesis pathway modules ('SUBSPEC': defined with substrate-specific enzymes whenever possible; 'BUCASYNOP': the 6-gene core operon for butyryl-CoA synthesis; 'BUTANOATE': the full set of enzymes with all potential alternative KOs). The first column indicates the genome and subsequent columns indicate the module completeness scores as computed from the KOfams annotated by each tool (with default parameters).

**Supplementary Table 3.** Top 10 most dissimilar modules (highest absolute difference) based on average completeness scores averaged across per family for anvi'o and MicrobeAnnotator. Results from MicrobeAnnotator were added for comparison.

**Supplementary Table 4.** Percent of modules per family whose completeness scores increased using anvi'o or MicrobeAnnotator.
